# Controlled Substrate Crossover from Cathode to Anode for Long-Term Autonomous Operation of Microbial Fuel Cells: A Transport–Reaction Modeling Study

**DOI:** 10.64898/2026.08.14.744300

**Authors:** Marcelo Gamboa Velasquez, Raul Gonzalo Meneses Sandoval, Jose Manuel Balderrama Perez, Maria Esther Medina Villafuerte, Jerry Luis Solis Valdivia

## Abstract

Microbial fuel cells (MFCs) have been widely investigated as decentralized bioelectrochemical systems capable of converting organic substrates into electricity. However, their long-term autonomous operation is constrained by substrate depletion in the anode compartment, leading to metabolic starvation of electroactive biofilms and a decline in power output. Conventional MFC design treats substrate crossover through the membrane separator as a parasitic loss that reduces coulombic efficiency. In this work, we propose a conceptual inversion of this paradigm by considering controlled cathodic-to-anodic substrate crossover as a passive mechanism to sustain basal microbial metabolism during periods of substrate scarcity. A transport–reaction framework is developed to quantify the balance between membrane-mediated substrate flux and microbial maintenance demand within the anode biofilm. Based on this balance, a dimensionless maintenance crossover Damköhler number (*Da_m_*) is introduced to define three operational regimes: starvation-dominated (*Da_m_ ≫* 1), balanced autonomous (*Da_m_ ≈* 1), and crossover-dominated (*Da_m_ ≪* 1). The framework integrates membrane transport theory with biofilm kinetics to evaluate the effects of separator properties, substrate gradients, and current-dependent electro-osmotic transport on system stability. Order-of-magnitude analysis indicates that achievable crossover fluxes span several orders of magnitude depending on separator characteristics, suggesting that membrane properties critically influence system behavior. This perspective reframes substrate crossover from a loss mechanism to a potential design variable, offering a conceptual tool for enhancing resilience and guiding separator selection in MFCs intended for long-duration, and low-maintenance operation.

**Highlights:**

- Controlled crossover can sustain microbial metabolism in MFCs
- Introduces maintenance crossover Damköhler number (*Da_m_*)
- Identifies regimes for autonomous and starvation operation
- Links membrane properties to long-term system stability
- Reframes crossover as a design variable, not only a loss

## 1. Introduction

### 1.1. Microbial fuel cells: principles and recent advances

Microbial fuel cells (MFCs) are a class of bioelectrochemical systems in which electroactive microorganisms oxidize organic substrates and transfer the released electrons to an anode, enabling direct conversion of biochemical energy into electrical current [1–3]. This process is mediated by extracellular electron transfer (EET) mechanisms, including direct electron transfer via conductive pili or outer membrane cytochromes, as well as mediated pathways involving soluble redox shuttles [4–7]. In dual-chamber configurations, anodic oxidation of soluble electron donors is coupled to cathodic reduction reactions—most commonly oxygen reduction—through an external electrical circuit and an ion-conducting separator that maintains electrochemical continuity while limiting bulk mixing of the two compartments [8–10]. The separator plays a critical role in sustaining ionic transport while modulating the exchange of chemical species between anodic and cathodic environments [10–12]. Over the past two decades, extensive research has led to substantial improvements in electrode materials, reactor architectures, and microbial enrichment strategies [2,13–15]. These advances have enabled laboratory-scale power densities approaching 1–3 W m*^−^*^2^ under optimized and continuously fed conditions [2,13,16,17]. As a result, MFCs have emerged as promising platforms for decentralized wastewater treatment, environmental monitoring, and low-power energy recovery [18–20], particularly in applications where conventional energy infrastructure is limited [21].

### 1.2. Operational limitations: substrate dependence and system fragility

Despite these technical developments, the transition of MFCs from controlled laboratory systems to robust field technologies remains constrained by fundamental operational limitations [2,13]. Among these, the dependence of electroactive biofilms on continuous substrate availability represents a critical bottleneck [22,23]. High-performing anodic communities rely on sustained access to soluble organic electron donors to maintain metabolic activity and electron flux [22–24]. When substrate supply is interrupted, current production typically declines rapidly—often by an order of magnitude within hours to days—reflecting the tight coupling between metabolic turnover and electrical output [25–27]. Prolonged starvation leads to metabolic downregulation, reduced expression of extracellular electron transfer pathways, and, in some cases, irreversible loss of electrogenic activity through cellular dormancy, community restructuring, or biomass decay [23,28]. Consequently, conventional MFC operation relies on continuous feeding regimes or frequent operator intervention to maintain stable performance [22]. This requirement is incompatible with many of the applications for which MFCs are most attractive, including remote environmental sensing, off-grid monitoring systems, sediment and benthic deployments, and wastewater treatment in resource-limited settings where long-term autonomous operation is essential [18,21].

### 1.3. Dominant design paradigm: suppression of crossover

Within the dominant design paradigm of MFC research, maximizing instantaneous power density and coulombic efficiency has been the primary objective [2,13]. To achieve these goals, system configurations are typically optimized to minimize internal losses associated with mass transport between compartments [2,29]. Two transport phenomena have received particular attention: oxygen diffusion from cathode to anode, which disrupts the anaerobic conditions required by many exoelectrogenic microorganisms [29,30], and crossover of soluble organic substrates from the catholyte to the anolyte, which reduces coulombic efficiency by enabling competing metabolic pathways or promoting cathodic fouling [31–33]. To suppress these effects, highly selective ion-exchange membranes—most commonly proton or cation exchange membranes such as Nafion—are widely employed due to their low permeability to organic solutes and high ionic conductivity [2,29]. While this approach is effective for maximizing performance under continuously fed laboratory conditions, it also functionally isolates the anodic compartment from potential passive substrate sources [34]. As a result, once external feeding ceases, the anodic biofilm experiences rapid substrate depletion, and the system transitions into a starvation-driven decline in electrochemical activity [25].

### 1.4. Evidence from resilient bioelectrochemical systems

In contrast to highly engineered laboratory configurations, several naturally occurring or minimally engineered bioelectrochemical systems demonstrate remarkable long-term stability despite limited substrate inputs [35]. Sediment-based and benthic MFCs, for example, can sustain low but persistent current production over months or even years through slow diffusive transport of organic compounds from surrounding sediments or aqueous environments [26,36,37]. In these systems, the anodic biofilm is not completely isolated but instead receives a continuous, low-level substrate supply mediated by passive mass transport processes [38]. Similarly, pilot-scale MFC systems operated under intermittent feeding conditions have shown that electroactive microbial communities can remain viable and capable of rapid reactivation following extended starvation periods [39,40]. These observations suggest that complete elimination of passive substrate transport may not always represent the optimal strategy, particularly in applications where longterm stability, resilience, and minimal maintenance are prioritized over peak instantaneous performance [35].

### 1.5. Conceptual reframing: crossover as a resource

Motivated by these observations, this work proposes a conceptual reframing of substrate crossover in dual-chamber MFCs. Rather than treating the passive transport of soluble organics across the separator exclusively as a parasitic loss mechanism, we examine the possibility that controlled cathodicto-anodic crossover can function as a passive substrate supply mechanism capable of sustaining basal metabolic activity under substrate-limited conditions [31,41]. In this framework, the catholyte acts as a substrate reservoir, while the separator regulates the diffusive and electro-osmotic transport of organic molecules toward the anode. We hypothesize that, under appropriate conditions, the magnitude of this transmembrane flux can be tuned such that it is sufficiently small to avoid significant efficiency losses, yet large enough to satisfy the maintenance metabolic requirements of the anodic biofilm. Similar transport–reaction balances have been observed in membrane bioreactors and electrochemical systems where coupling between flux and reaction stabilizes system performance [29,42].

### 1.6. Dimensionless framework: maintenance crossover Damköhler number

To quantify the relationship between membrane-mediated substrate transport and microbial metabolic demand, we introduce a dimensionless parameter termed the maintenance crossover Damköhler number (*Da_m_*). In classical reaction–transport systems, Damköhler numbers compare characteristic reaction rates to transport rates, providing insight into whether system behavior is reaction-limited or transport-limited [43,44]. In the present context, *Da_m_* is defined as:

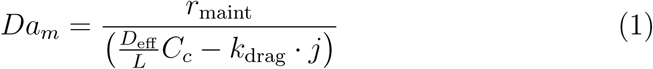

where *r*_maint_ represents the areal maintenance substrate consumption rate, *D*_eff_ is the effective diffusivity of the substrate within the separator material, *L* is the separator thickness, *C_c_* is the bulk substrate concentration in the catholyte, *k*_drag_ is an effective electro-osmotic drag coefficient describing current-induced solvent transport, and *j* is the operating current density. This formulation captures the balance between biofilm metabolic demand and the maximum passive substrate supply achievable through coupled diffusive and electro-osmotic transport mechanisms [29,30].

### 1.7. Regime interpretation and design implications

When *Da_m_ ≪* 1, passive transport exceeds maintenance demand, potentially leading to substrate accumulation and diversion of metabolism toward non-electrogenic pathways, thereby reducing system efficiency [32,33]. Conversely, when *Da_m_ ≫* 1, transport is insufficient to sustain even basal metabolic requirements, resulting in progressive biofilm starvation and loss of electrochemical activity [25–27]. Between these limits lies a balanced regime (*Da_m_ ≈* 1), in which passive crossover approximately matches microbial maintenance requirements. In this regime, the anodic biofilm can theoretically sustain baseline metabolic activity without continuous external feeding. Furthermore, electro-osmotic transport coupled to current generation may introduce a weak stabilizing feedback between metabolic activity and substrate availability, as observed in related electrochemical transport systems [29,42]. By framing separator transport capacity and microbial metabolic demand within a unified dimensionless framework, this analysis identifies membrane permeability, separator thickness, catholyte composition, and operating current density as coupled design variables that jointly determine long-term system stability [2,29]. Importantly, this perspective shifts the design objective from strict minimization of crossover flux toward optimization of the balance between transport supply and biological demand, enabling new strategies for resilient and autonomous MFC operation.

### 1.8. Scope and structure of this work

The remainder of this study develops the theoretical basis of the proposed framework. Section 2 reviews the physical and literature foundations of substrate crossover in microbial fuel cells. Section 3 presents the transport–reaction model for membrane-mediated substrate supply and introduces the maintenance crossover Damköhler number. Section 4 develops an engineering design map for autonomous bioelectrochemical systems and examines the resulting operational regimes. Section 5 discusses reaction–transport coupling and broader theoretical implications. Section 6 presents the conclusions.

## 2. Physical and Literature Foundations of Substrate Crossover in Microbial Fuel Cells

### 2.1. Coupled transport–reaction processes in dual-chamber systems

Dual-chamber microbial fuel cells (MFCs) are inherently coupled transport– reaction systems in which electrochemical processes, microbial metabolism, and mass transfer occur across spatially segregated domains linked by an ion-conducting separator [1,2]. The anodic compartment hosts electroactive biofilms that oxidize dissolved organic substrates, generating electrons that are transferred to the anode through extracellular electron transfer pathways. These electrons flow through an external circuit to the cathode, where they are consumed by a terminal reduction reaction, typically oxygen reduction. The closure of the electrical circuit requires ionic transport through the separator to maintain electroneutrality [45]. This transport is mediated by mobile charge carriers (e.g., protons, cations, or anions depending on membrane type), but it is intrinsically coupled to the transport of neutral and charged solutes as well as solvent [46]. Consequently, the separator does not function as an ideal selective barrier but rather as a heterogeneous transport medium in which multiple species are exchanged under coupled driving forces [12, 47]. From a continuum perspective, system behavior is governed by the interaction between: (i) electrochemical reaction rates at the electrodes, (ii) microbial substrate consumption within the anodic biofilm, and (iii) multicomponent transport across the separator and adjacent boundary layers [48–50]. These processes operate on comparable spatial scales and are dynamically linked through concentration gradients, electric fields, and current-dependent transport phenomena [51]. As a result, the separator constitutes a critical control volume in which small variations in material properties or operating conditions can significantly alter overall system behavior [52]. Key separator characteristics influencing transport include thickness *L*, porosity *ε*, tortuosity *τ* , fixed charge density, and hydration state [53, 54]. These parameters determine not only ionic conductivity and ohmic resistance but also permeability to dissolved species and the magnitude of solvent fluxes [53]. Accurate description of MFC operation therefore requires explicit consideration of the mechanisms governing transmembrane transport.

### 2.2. Mechanisms of solute transport across separators

Transport of dissolved species in membrane-separated electrochemical systems is fundamentally described by the Nernst–Planck framework, which accounts for the superposition of diffusive, migrational, and convective fluxes [55, 56]. The molar flux *J_i_* of species *i* can be expressed as:

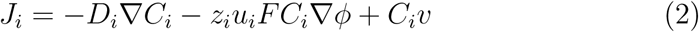

where *D_i_*is the molecular diffusivity, *C_i_*the concentration, *z_i_*the ionic valence, *u_i_*the ionic mobility, *F* the Faraday constant, *ϕ* the electric potential, and *v* the solvent velocity.

#### 2.2.1. Effective transport in porous and ion-exchange media

In practical MFC separators, transport occurs within a complex microstructure characterized by pores, channels, or hydrated polymer domains [53, 57]. The intrinsic transport properties must therefore be reformulated in terms of effective parameters [58]. For diffusive transport, this yields:

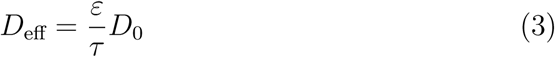

where *D*_0_ is the bulk diffusivity, *ε* the porosity (or water volume fraction in polymer membranes), and *τ* the tortuosity factor accounting for increased path length and geometric constraints. In ion-exchange membranes, additional effects arise from fixed charges within the polymer matrix [59, 60]. These charges generate Donnan potentials at the membrane–solution interface, leading to selective exclusion of co-ions and modification of ion partitioning [59]. For weakly charged or neutral organic substrates, this typically results in reduced permeability relative to bulk solution, although the extent of exclusion depends on molecular size, polarity, and membrane hydration [61].

#### 2.2.2. Diffusion-dominated transport of organic substrates

For the low-molecular-weight organic compounds commonly used in MFCs (e.g., acetate, lactate, propionate), transmembrane transport is generally dominated by diffusion [53]. Although such species may exist in partially dissociated form under operating conditions, their effective migration through ion-exchange membranes is strongly limited by electrostatic exclusion and low mobility within the membrane phase [62]. Under these conditions, the substrate flux across a separator of thickness *L* can be approximated using a one-dimensional Fickian description:

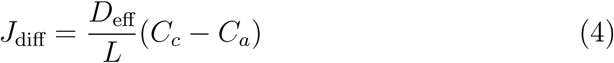

where *C_c_* and *C_a_* denote the substrate concentrations at the catholyte bulk and the anode–membrane interface, respectively. This formulation assumes quasi-steady transport and negligible accumulation within the membrane phase, conditions that are typically satisfied due to the small characteristic thickness of separators relative to reactor dimensions [63].

#### 2.2.3. Electro-osmotic and current-coupled transport

In addition to diffusion, solvent-mediated transport can contribute to solute flux through electro-osmotic effects [64, 65]. In hydrated membranes, migration of charge carriers (e.g., protons in proton-exchange membranes) induces a net flux of water molecules through viscous coupling at the molecular scale [64]. This solvent flux is proportional to current density and can entrain dissolved species present within the membrane [65]. The resulting convective contribution to solute transport can be expressed as:

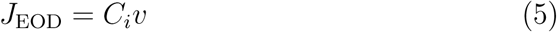

with solvent velocity approximated by:

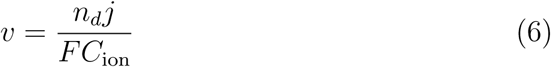

where *n_d_* is the electro-osmotic drag coefficient (number of solvent molecules transported per charge carrier), *j* the current density, and *C*_ion_ the concentration of mobile ions within the membrane phase. For modeling purposes, this contribution is often represented in lumped form as:

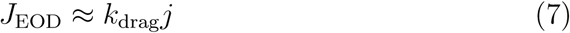

where *k*_drag_ incorporates membrane structure, hydration, and solute partitioning effects. Although electro-osmotic transport is typically smaller than diffusive fluxes under dilute conditions, it introduces a direct coupling between electrochemical activity and mass transport [66]. Depending on current direction and membrane properties, this mechanism can either enhance or oppose diffusive substrate flux, thereby influencing net transmembrane transport under dynamic operating conditions [66].

### 2.3. Impacts of substrate crossover on electrochemical and biological processes

Transmembrane transport of organic substrates affects MFC performance through multiple physicochemical and biological pathways [31, 67]. From an electrochemical standpoint, the redistribution of substrates between compartments alters the spatial distribution of electron donors and acceptors. Substrates that bypass anodic oxidation represent a loss of recoverable chemical energy and can contribute to undesired reactions at the cathode, including microbial growth, catalytic side reactions, or fouling processes that increase interfacial resistance [68,69]. From a biological perspective, the presence of alternative electron donors or acceptors in either compartment can shift metabolic pathways within microbial communities. In the anode, competition between exoelectrogenic respiration and alternative pathways (e.g., fermentation or methanogenesis) can reduce the fraction of electrons recovered as current [67, 70]. In the cathode, substrate availability may support heterotrophic growth, leading to biofilm formation and transport limitations [31]. In addition, solute transport is coupled to ionic fluxes and can influence pH gradients, buffering capacity, and the distribution of inorganic species. These effects may contribute to precipitation, membrane fouling, and longterm degradation of separator performance [71, 72]. The magnitude and direction of these impacts depend on the relative rates of transport and reaction, highlighting the need for quantitative frameworks capable of resolving their interplay.

### 2.4. Separator materials and transport–selectivity relationships

A wide range of separator materials has been investigated in MFC systems, including dense ion-exchange membranes, porous polymer separators, ceramics, and composite structures. These materials exhibit distinct transport characteristics that reflect trade-offs between ionic conductivity, selectivity, and permeability. Dense ion-exchange membranes, such as perfluorosulfonic acid polymers, are characterized by high ionic conductivity and strong electrostatic exclusion of co-ions. Their hydrated nanophase-separated structure enables efficient proton transport while limiting the passage of larger or weakly charged organic molecules. However, finite permeability remains due to diffusion through aqueous domains and imperfections in membrane structure [73, 74]. Porous separators, including nonwoven fabrics and ceramics, exhibit higher permeability to dissolved species due to their open pore structure. In these materials, transport is governed primarily by diffusion and convection within interconnected pores, with limited selectivity. While such separators reduce ohmic resistance and can enhance mass transfer, they also permit greater crossover of substrates and other solutes [75, 76]. Composite and functionalized membranes seek to balance these competing effects by combining structural control with chemical selectivity. Modifications such as surface functionalization, incorporation of inorganic fillers, or multilayer architectures can alter effective diffusivity, charge selectivity, and hydration behavior [77, 78, 79]. From a transport perspective, separator performance can be characterized by an effective permeability *P* = *D*_eff_*/L*, which directly determines diffusive flux under a given concentration gradient [71, 76, 80]. Variations in permeability across different materials span several orders of magnitude, implying that separator selection can fundamentally alter the extent of transmembrane transport and its impact on system behavior [76, 81].

### 2.5. Limitations of existing modeling approaches

Most modeling frameworks for MFCs have been developed under conditions of continuous substrate supply and are therefore formulated to describe steady-state operation. These models typically couple Monod-type kinetics for microbial substrate utilization with charge transport, electrode kinetics, and diffusion within biofilms and boundary layers [82]. Within such formulations, transmembrane substrate transport is often treated as a secondary loss term or neglected altogether [83]. This approach is appropriate when external substrate supply dominates system dynamics, but it becomes inadequate when transport across the separator contributes significantly to substrate availability. Furthermore, conventional dimensionless analysis in reaction engineering relies on characteristic timescales associated with flow reactors or diffusion-reaction systems with well-defined boundary conditions [84, 85]. In closed or intermittently fed MFCs, these timescales are not directly applicable, as system behavior may instead be governed by the balance between slow transport processes and minimal metabolic demand. As a result, existing models provide limited insight into regimes where passive transport controls substrate availability, and they do not offer general criteria for assessing whether such transport is sufficient to sustain microbial activity.

### 2.6. Flux-based description of transport–reaction coupling

A physically consistent description of separator-mediated substrate supply requires formulation in terms of flux balances rather than residence times [86]. In this framework, the relevant comparison is between: the maximum substrate flux that can be delivered to the anode via transmembrane transport, and the rate at which the anodic biofilm consumes substrate to sustain metabolic activity. The transport-limited flux is determined by the combined contributions of diffusion and current-coupled convection:

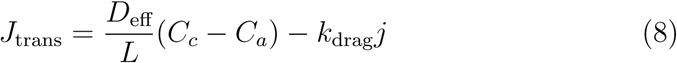

while the consumption rate depends on biofilm properties, microbial physiology, and local substrate concentration. Formulating the problem in this manner enables direct evaluation of whether transport processes can supply substrate at a rate commensurate with biological demand. This flux-based perspective provides the necessary foundation for defining dimensionless parameters that characterize system behavior across different operating regimes. The following section builds upon this formulation to develop a transport–reaction model that quantitatively links separator properties, operating conditions, and microbial kinetics, leading to the definition of a dimensionless criterion for assessing the role of substrate crossover in sustaining anodic activity.

## 3. Transport–Reaction Model for Membrane-Mediated Substrate Supply

### 3.1. Modeling Scope and Physical Assumptions (Revised)

To evaluate whether passive substrate crossover can sustain microbial activity under substrate-limited conditions, a transport–reaction framework is formulated for a dual-chamber microbial fuel cell (MFC), as schematically illustrated in Figure 1. The system consists of anodic and cathodic compartments separated by a membrane or porous separator of thickness *L* (m) and area *A* (m²). The catholyte contains a dissolved organic substrate at bulk concentration *C_c_* (mol m*^−^*^3^), while the anode hosts an electroactive biofilm that consumes substrate at the electrode surface. Under the well-mixed assumption, the interfacial concentration is approximated by the bulk anolyte concentration *C_a_* (mol m*^−^*^3^). The objective is to determine whether passive transmembrane transport can supply substrate at a rate sufficient to balance microbial maintenance demand during starvation. To retain analytical tractability while preserving the dominant transport–reaction physics, the following assumptions are adopted:

- Neutral substrate species: electromigration is neglected relative to diffusion.
- One-dimensional transport across the separator thickness.
- No reaction within the separator; consumption is confined to the anode.
- Finite maintenance current: the system operates at a low but non-zero current density *j* (A m*^−^*^2^).
- No pressure-driven flow.
- Negligible external mass transfer resistance (well-mixed bulk phases).
- Isothermal conditions with constant transport coefficients.

**Figure 1:**
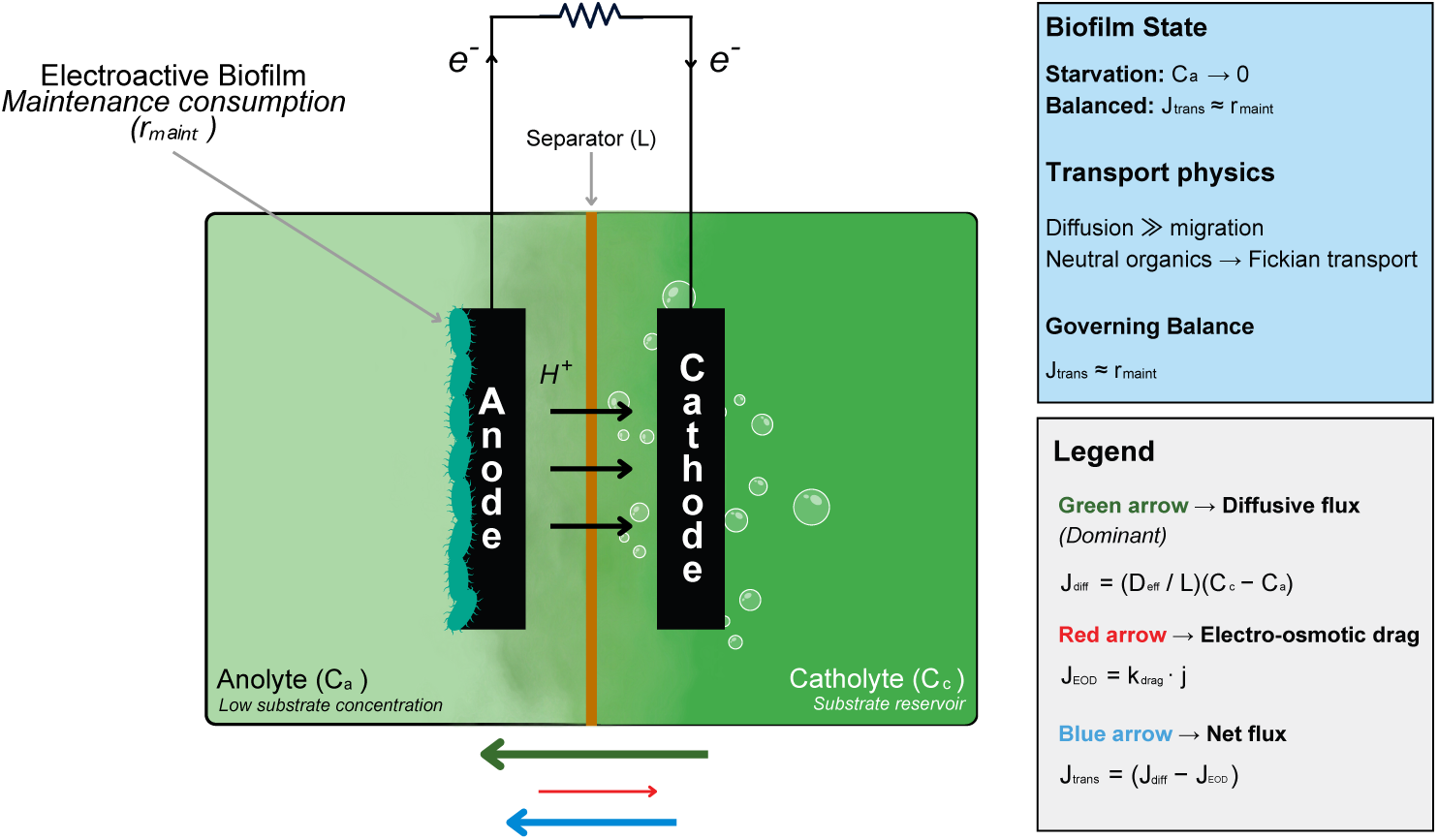
Schematic illustration of a dual-chamber microbial fuel cell with controlled cathodic-to-anodic substrate crossover as a passive maintenance supply mechanism. The net transmembrane substrate flux 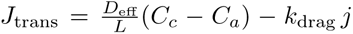 balances against the maintenance consumption rate *r*_maint_ of the anodic biofilm in the starvation limit (*C_a_ →* 0). This conceptual reframing shifts substrate crossover from a parasitic loss to a tunable passive supply mechanism for sustaining long-term biofilm viability.

All consumption rates are expressed per unit membrane area (mol m*^−^*^2^ s*^−^*^1^) to ensure consistency with transmembrane fluxes.

### 3.2. Transmembrane Substrate Flux

Flux is defined as positive in the cathode-to-anode direction.

#### 3.2.1. Diffusive transport

Under quasi-steady, one-dimensional conditions, substrate transport across the separator is dominated by diffusion [76, 81, 87].

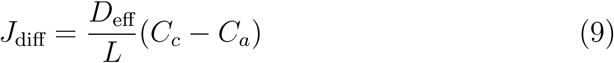

where *D*_eff_ is the effective diffusivity accounting for membrane microstructure, including porosity, tortuosity, and solute partitioning.

#### 3.2.2. Electro-osmotic contribution

Ionic current through hydrated membranes induces solvent transport via electro-osmotic drag. This solvent flux entrains dissolved species and introduces a convective contribution to substrate transport [88]. A lumped representation is adopted:

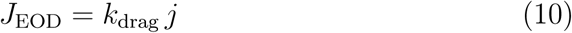

where *k*_drag_ (mol m*^−^*^2^ s*^−^*^1^ per A m*^−^*^2^) is an effective electro-osmotic drag coefficient that incorporates solvent flux coupling, membrane hydration, and solute partitioning effects. In cation-exchange membranes typical of MFCs, electro-osmotic flow is directed toward the cathode and therefore opposes diffusive transport toward the anode [9, 71].

#### 3.2.3. Net substrate flux

The total transmembrane substrate flux is:

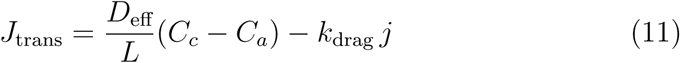

This expression defines the upper bound of substrate delivery under transport-limited conditions, as it neglects any kinetic limitations within the biofilm.

### 3.3. Biofilm Substrate Consumption

Substrate consumption by the anodic biofilm is described using Monod-type kinetics:

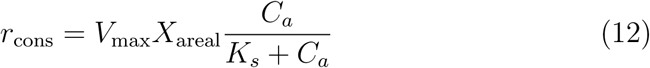

where *V*_max_ is the maximum specific uptake rate, *X*_areal_ the areal biomass density, and *K_s_* the half-saturation constant. Under substrate-limited conditions (*C_a_ ≪ K_s_*):

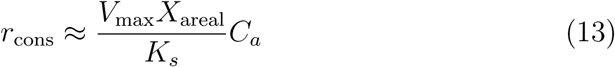

In addition to growth-associated consumption, electroactive biofilms require a baseline metabolic flux to sustain cellular maintenance [89]. This is represented as a minimum areal consumption rate *r*_maint_ (mol m*^−^*^2^ s*^−^*^1^). The maintenance consumption rate can be related to a baseline current density through Faradaic conversion [90, 91]:

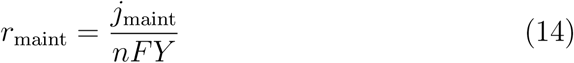

where *n* is the number of electrons transferred per mole of substrate, *F* is Faraday’s constant (96485 C mol*^−^*^1^), and *Y* is an effective yield coefficient relating substrate consumption to current generation.

### 3.4. Maintenance Crossover Damköhler Number

In the starvation limit (*C_a_ →* 0), the driving force for transport is maximized [90]. The maximum achievable substrate supply is defined as:

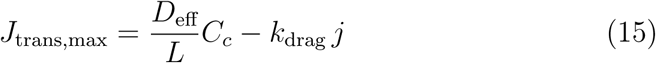

This expression accounts for both diffusive and electro-osmotic contributions to transport. A dimensionless criterion is obtained by comparing maintenance demand to this transport-limited supply:

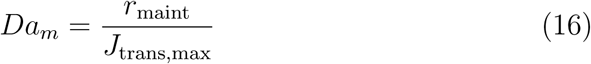

This parameter quantifies the balance between microbial maintenance requirements and the maximum transport-limited substrate flux.

### 3.5. Operational Regimes

The magnitude of *Da_m_*defines three distinct regimes:

- Transport-limited starvation (*Da_m_ ≫* 1) Passive transport is insufficient to meet maintenance demand, leading to substrate depletion, metabolic downregulation, and decay of electrochemical activity.
- Balanced autonomous regime (*Da_m_ ≈* 1) Transport approximately matches maintenance demand, sustaining a finite substrate concentration and enabling long-term metabolic activity without external feeding.
- Excess crossover (*Da_m_ ≪* 1) Transport exceeds metabolic demand, leading to substrate accumulation and potential diversion toward non-electrogenic pathways, reducing coulombic efficiency.

### 3.6. Dynamic Mass Balance in the Anode

The transient substrate concentration in the anode satisfies:

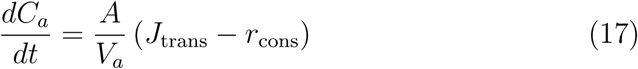

Substituting the flux expression:

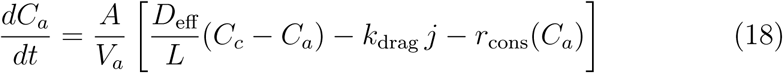

Steady state is achieved when:

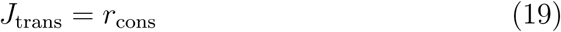

corresponding to the dynamically realized balance associated with *Da_m_ ≈* 1.

### 3.7. Governing Parameters and Design Implications

The model demonstrates that passive substrate supply is governed by a small set of coupled parameters:

- separator transport capacity *D*_eff_*/L*,
- catholyte substrate concentration *C_c_*,
- electro-osmotic coupling *k*_drag_ *j*,
- biofilm maintenance demand *r*_maint_.

These variables define a design space in which membrane properties, operating conditions, and microbial characteristics jointly determine system behavior. This framework shows that, rather than being strictly detrimental, substrate crossover can be tuned to achieve a balance between transport supply and biological demand. By adjusting separator permeability and operating current density, it is possible to approach the balanced regime in which passive transport sustains microbial maintenance without significant efficiency loss. This formulation establishes a physically consistent basis for evaluating the feasibility of passive substrate supply and provides the foundation for the regime analysis developed in the following section.

## 4. Engineering Design Map for Autonomous Bioelectrochemical Systems

The transport–reaction framework developed in Section 3 establishes that microbial viability under substrate-limited conditions is governed by the maintenance crossover Damköhler number:

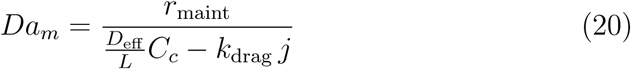

which compares microbial maintenance demand to the maximum transport-limited substrate flux from the cathode to the anode. In the present section, this theoretical formulation is translated into quantitative engineering constraints by mapping the governing parameters onto realistic ranges encountered in microbial fuel cells (MFCs). The objective is to identify the conditions under which passive substrate crossover can sustain anodic biofilm activity over extended time scales.

### 4.1. Representative Parameter Ranges

The governing parameters span multiple orders of magnitude depending on separator material, reactor design, and microbial community. Representative ranges consistent with reported MFC systems are [90, 91, 92]:

- Bulk substrate diffusivity: *D*_0_ *∼* (0.7 *−* 1.2) *×* 10*^−^*^9^ m^2^ s*^−^*^1^
- Effective diffusivity in separator: *D*_eff_ *∼* 5 *×* 10*^−^*^11^ *−* 4 *×* 10*^−^*^10^ m^2^ s*^−^*^1^
- Separator thickness: *L ∼* 10*^−^*^4^ *−* 10*^−^*^3^ m
- Catholyte substrate concentration: *C_c_ ∼* 10 *−* 200 mol m*^−^*^3^
- Maintenance current density: *j*_maint_ *∼* 0.01 *−* 0.1 A m*^−^*^2^
- Maintenance substrate consumption rate: *r*_maint_ *∼* 10*^−^*^7^ *−* 10*^−^*^5^ mol _m_*−*2 _s_*−*1

These ranges highlight that separator-controlled transport properties, particularly *D*_eff_ and *L*, can vary over several orders of magnitude and therefore act as primary determinants of system behavior [90, 92]. To facilitate quantitative interpretation and reproducibility, representative parameter values and derived transport quantities are summarized in Table 1. The values reported in Table 1 provide the quantitative basis for the order-of-magnitude analysis and regime classification developed in the following sections.

**Table 1:** Representative ranges of key parameters in microbial fuel cells and corresponding order-of-magnitude estimates for diffusive and electro-osmotic substrate fluxes. summarizes parameter ranges drawn from MFC literature. Fluxes are calculated in the starvation limit (*Ca →* 0). Diffusion dominates by 2–3 orders of magnitude under typical conditions, while separator properties (*D*_eff_ and *L*) shift *Da_m_* across regimes.

| (a) Primary parameters |  |  |  |  |
| --- | --- | --- | --- | --- |
| Parameter | Symbol | Typical range / value | Units | Notes / Literature basis |
| Bulk diffusivity | $D_0$ | $(0.7\text{--}1.2) \times 10^{-9}$ | $\text{m}^2 \text{s}^{-1}$ | Small organic molecules (e.g., acetate) in water |
| Effective diffusivity (low permeability) | $D_{\text{eff,low}}$ | $1 \times 10^{-11}$ | $\text{m}^2 \text{s}^{-1}$ | Dense ion-exchange membranes |
| Effective diffusivity (high permeability) | $D_{\text{eff,high}}$ | $5 \times 10^{-10}$ | $\text{m}^2 \text{s}^{-1}$ | Porous separators / hydrated structures |
| Separator thickness (reference value) | $L$ | $5 \times 10^{-4}$ | m | Representative mid-range value used for derived quantities |
| Catholyte substrate concentration | $C_c$ | 10–200 | mol $\text{m}^{-3}$ | Representative dissolved organic concentration |
| Maintenance current density | $j_{\text{maint}}$ | 0.01–0.1 | $\text{A m}^{-2}$ | Basal metabolic activity |
| Maintenance consumption rate | $r_{\text{maint}}$ | $1 \times 10^{-6}$ | mol $\text{m}^{-2} \text{s}^{-1}$ | Representative value for regime analysis |
| Electro-osmotic drag coefficient | $k_{\text{drag}}$ | $1 \times 10^{-6}$ | mol $\text{m}^{-2} \text{s}^{-1} (\text{A m}^{-2})^{-1}$ | Lumped transport coefficient |
| Operating current density (reference) | $j$ | 0.05 | $\text{A m}^{-2}$ | Representative operating condition |
| (b) Derived transport quantities (order-of-magnitude analysis) |  |  |  |  |
| (Calculated using $L = 5 \times 10^{-4} \text{ m}$ and $C_c = 100 \text{ mol m}^{-3}$) | | | | |
| Quantity | Expression | Low permeability | High permeability | Units |
| Transport coefficient | $D_{\text{eff}}/L$ | $2 \times 10^{-8}$ | $1 \times 10^{-6}$ | $\text{m s}^{-1}$ |
| Maximum diffusive flux | $J_{\text{diff,max}} = (D_{\text{eff}}/L) C_c$ | $2 \times 10^{-6}$ | $1 \times 10^{-4}$ | mol $\text{m}^{-2} \text{s}^{-1}$ |
| Electro-osmotic flux | $J_{\text{EOD}} = k_{\text{drag}} j$ | $5 \times 10^{-8}$ | $5 \times 10^{-8}$ | mol $\text{m}^{-2} \text{s}^{-1}$ |
| Net transport flux | $J_{\text{trans,max}} = J_{\text{diff,max}} - J_{\text{EOD}}$ | $\approx 2 \times 10^{-6}$ | $\approx 1 \times 10^{-4}$ | mol $\text{m}^{-2} \text{s}^{-1}$ |
| Damköhler number | $Da_m = r_{\text{maint}}/J_{\text{trans,max}}$ | $\approx 0.5$ | $\approx 0.01$ | – |

### 4.2. Maximum Transport-Limited Substrate Flux

In the starvation limit (*C_a_ →* 0), the maximum achievable substrate flux is:

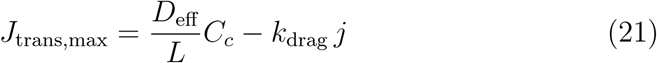

Using representative values [29, 93, 94]: *D*_eff_ = 5*×*10*^−^*^10^ m^2^ s*^−^*^1^, *L* = 5*×*10*^−^*^4^ m, *C_c_* = 100 mol m*^−^*^3^ yields:

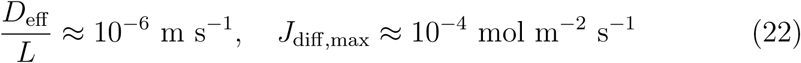

For a typical current density [95, 96] *j* = 0.05 A m*^−^*^2^ and *k*_drag_ *≈* 10*^−^*^6^ mol _m_*−*2 _s_*−*1 _(A m_*−*2_)_*−*1:

Thus:

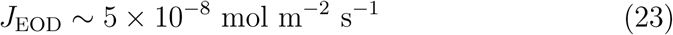

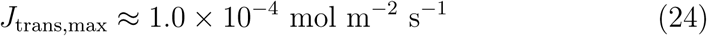

indicating that diffusion dominates transport by two to three orders of magnitude under typical operating conditions, while electro-osmotic effects remain secondary but non-negligible at higher current densities. These order-of-magnitude estimates establish the transport capacity that defines the upper bound for substrate supply and form the basis for the regime classification developed in the following section.

### 4.3. Regime Classification via Damköhler Number

For a representative maintenance demand:

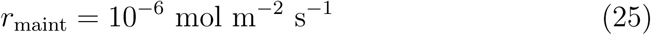

the corresponding Damköhler number is:

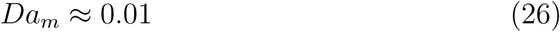

which lies in the crossover-dominated regime. For a low-permeability separator (*D*_eff_ = 1 *×* 10*^−^*^11^ m^2^ s*^−^*^1^):

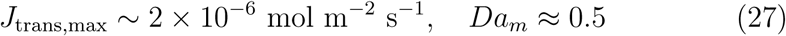

This transition illustrates that separator properties alone can shift system behavior across regimes without modification of microbial kinetics, emphasizing the dominant role of membrane-controlled transport. The resulting regime structure can be visualized as an engineering design map, as shown in Figure 2. This representation maps the dependence of the maintenance crossover Damköhler number on separator transport capacity (*D*_eff_ */L*) and catholyte concentration (*C_c_*), providing a direct link between the dimensionless formulation and practical design variables.

**Figure 2:**
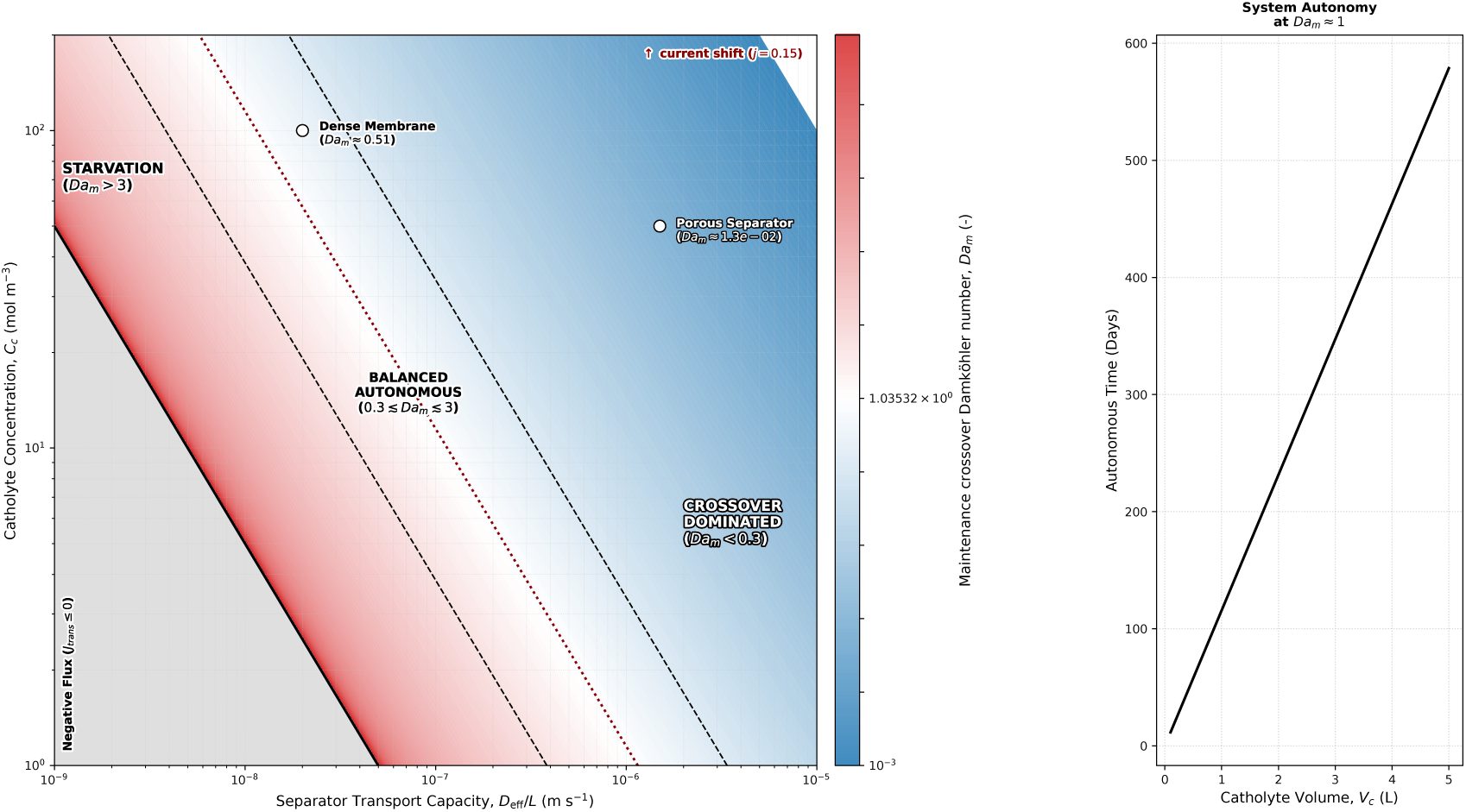
Engineering design map showing the three operational regimes defined by the maintenance crossover Damköhler number *Da_m_* = *r*_maint_*/* [(*D*_eff_ */L*)*C_c_ − k*_drag_*j*]. The balanced autonomous regime (0.3 ≲ *Da_m_* ≲ 3) enables passive substrate supply sufficient for maintenance metabolism without external feeding. Vertical dashed lines illustrate regime transitions caused by changes in separator thickness *L* (or equivalently transport capacity), while the weak effect of current density *j* through electro-osmotic drag is also shown.

### 4.4. Design Window for Autonomous Operation

As shown in Figure 2, the autonomous operating window corresponds to a bounded region in the parameter space defined by separator transport capacity and catholyte conditions. Transitions across this window reflect shifts between transport-limited and crossover-dominated behavior. Sustained autonomous operation requires a balance between transport supply and maintenance demand. This defines a bounded operational window:

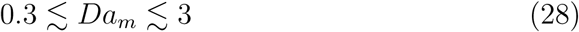

which can be rewritten as:

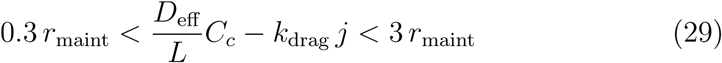

Within this window:

- substrate supply is sufficient to sustain maintenance metabolism, and
- excessive crossover leading to efficiency losses is avoided.

This inequality provides a direct and practical design criterion linking separator properties, operating conditions, and microbial demand, as visualized in the regime map presented in Figure 2.

### 4.5. Sensitivity to Separator Thickness

The diffusive contribution scales inversely with separator thickness:

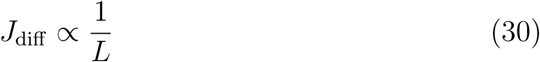

As a result, modest variations in thickness can induce regime transitions. For example:

- *L ≈* 0.5 mm: crossover-dominated or balanced regime
- *L ≈* 2 mm: transport-limited regime

This sensitivity explains why thinner or moderately permeable separators often exhibit improved long-term stability compared to dense ion-exchange membranes designed to minimize crossover [52, 53].

### 4.6. Catholyte Reservoir and Autonomy Time

The catholyte functions as a finite substrate reservoir:

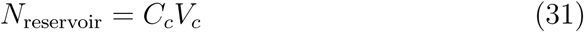

The characteristic autonomy time is:

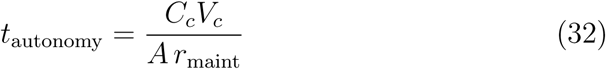

For representative values [18, 93, 97]: *C_c_* = 100 mol m*^−^*^3^, *V_c_* = 1 L, *A* = 0.01 m^2^, *r*_maint_ = 10*^−^*^6^ mol m*^−^*^2^ s*^−^*^1^ one obtains:

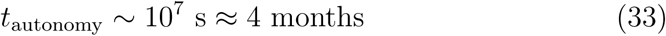

This scaling demonstrates that long-duration operation can be achieved through appropriate reservoir sizing, even in the absence of continuous feeding.

### 4.7. Current-Coupled Feedback Mechanism

Electro-osmotic transport introduces a weak coupling between current and substrate flux:

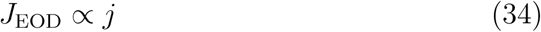

An increase in metabolic activity leads to higher current, which slightly reduces net substrate influx. This creates a stabilizing negative feedback loop: increased activity *→* higher current *→* reduced net transport *→* moderation of activity. Although secondary to diffusion under most conditions, this mechanism contributes to dynamic stability in the balanced regime.

### 4.8. Sensitivity and Uncertainty Analysis

The boundaries of the autonomous operating window are subject to uncertainty due to variability in key parameters:

- *D*_eff_ : sensitive to membrane microstructure, hydration, and fouling [24, 98]
- *r*_maint_: dependent on microbial community composition and physiological state [97, 99, 100]
- *k*_drag_: influenced by membrane type and ionic environment [101] A first-order sensitivity analysis shows:

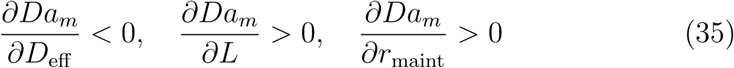

indicating that increases in separator permeability decrease *Da_m_*, while increases in thickness or maintenance demand increase it. These dependencies confirm that separator transport properties and microbial maintenance requirements are the dominant sources of uncertainty, as quantified by the first-order sensitivity coefficients and design boundaries summarized in Table 2.

**Table 2:**
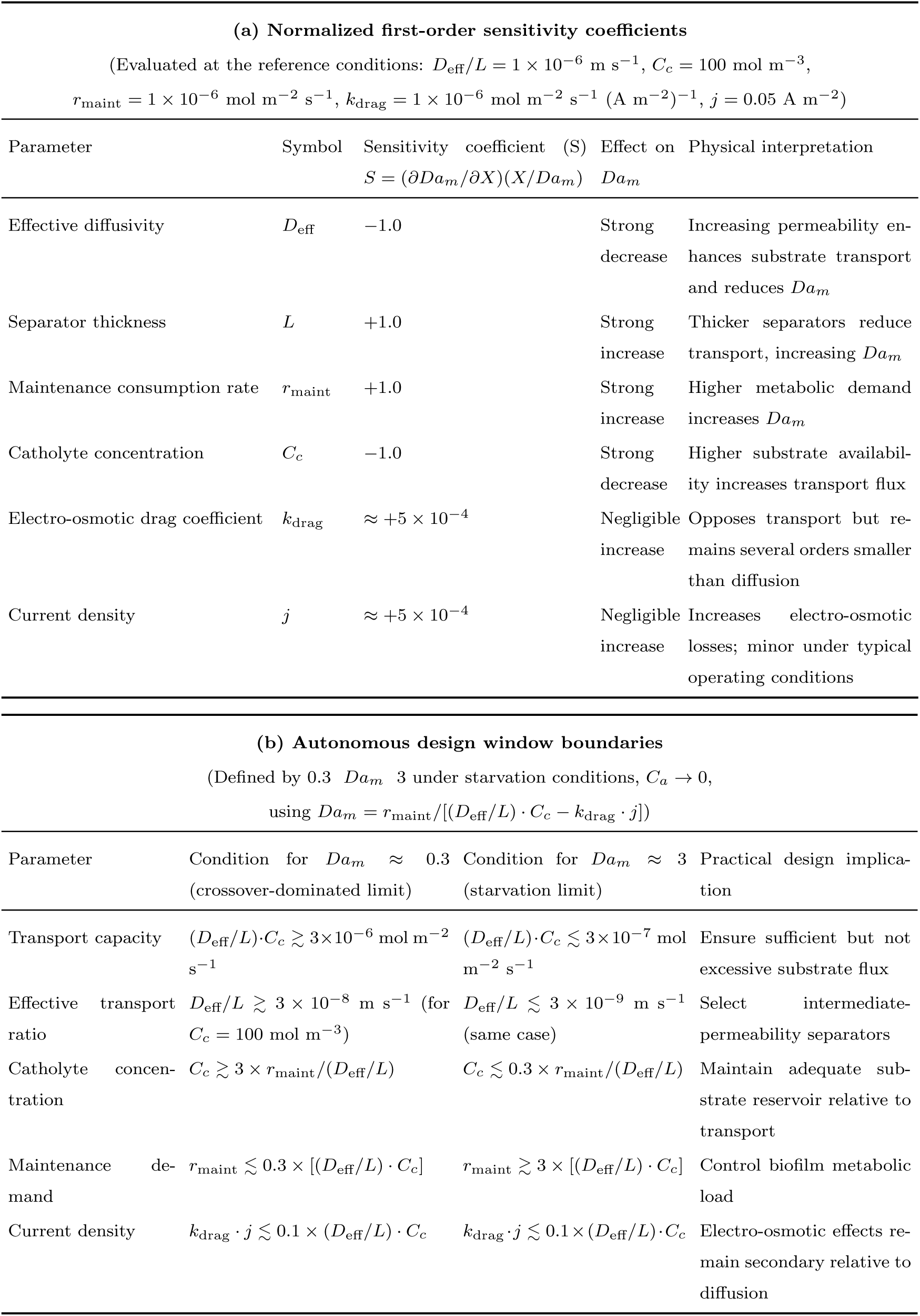
First-order sensitivity coefficients and boundaries of the autonomous design window (0.3 *Da_m_* 3) for representative microbial fuel cell conditions. Sensitivity analysis confirming that separator transport capacity (*D*_eff_*/L*) and maintenance demand *r*_maint_ dominate system behavior. The autonomous design window (0.3 *Da_m_* 3) identifies conditions under which passive substrate crossover sustains biofilm maintenance while minimizing efficiency losses. All sensitivities and boundary conditions are derived directly from the governing Damköhler expression under starvation conditions.

### 4.9. Design Implications for Autonomous MFCs

The engineering design map, supported by the sensitivity structure summarized in Table 2, yields several key principles:

- **Permeability optimization rather than minimization** Separator properties should be tuned to achieve *Da_m_ ≈* 1, rather than minimizing crossover indiscriminately.
- **Moderate-permeability separators as optimal solutions** Porous or semi-permeable materials can outperform highly selective membranes in long-term autonomous operation.
- **Catholyte as an engineered subsystem** The catholyte should be designed as a controlled substrate reservoir rather than treated as a passive compartment.
- **Coupled parameter design** Separator thickness, effective diffusivity, current density, and substrate concentration must be co-optimized within a unified framework.

### 4.10. Transition Toward Design-Oriented Frameworks

The results presented in this section establish a quantitative bridge between transport theory and reactor design. The analysis demonstrates that substrate crossover, traditionally treated as a parasitic loss, can instead be harnessed as a controllable transport pathway that stabilizes microbial metabolism under substrate-limited conditions. This shift from suppression to optimization defines a new design paradigm for microbial fuel cells intended for long-duration, low-maintenance operation.

## 5. Discussion: Reaction–Transport Coupling and Theoretical Implications

### 5.1. Reaction–transport perspective on maintenance-limited operation

The analysis developed in this work positions microbial fuel cells within the broader framework of reaction–transport systems, in which system behavior is governed by the balance between interfacial reaction rates and transport-limited supply. Unlike conventional catalytic or biological reactors, however, the relevant metabolic demand under substrate-limited conditions is not associated with growth or bulk conversion, but with the maintenance requirements of electroactive biofilms. This distinction fundamentally alters the governing design criterion. Rather than comparing reaction rates to residence times or bulk transport rates, the system is more appropriately described by a balance between maximum interfacial transport capacity and minimum metabolic demand. The resulting formulation, expressed as a ratio of areal fluxes at the anode–membrane interface, provides a physically consistent description that remains valid in the absence of well-defined hydrodynamic scales. This is particularly relevant for systems operating under intermittent feeding or quasi-closed conditions, where traditional reactor metrics lose predictive value. From this perspective, the transition between operational regimes is controlled by whether membrane-mediated transport can sustain a non-zero substrate concentration at the biofilm interface. The existence of a regime in which transport and maintenance demand are of comparable magnitude implies that long-term metabolic activity can be sustained without continuous external substrate supply.

### 5.2. Crossover as a controlled transport pathway

The results support a reinterpretation of substrate crossover as a functional transport pathway rather than an exclusively parasitic process. When separator permeability and catholyte composition are appropriately selected, transmembrane transport can provide a continuous, low-level substrate flux that stabilizes microbial metabolism during starvation. Under these conditions, the catholyte effectively acts as a distributed substrate reservoir, while the separator regulates delivery through its transport properties. This mechanism is analogous to diffusion-limited feeding in natural and engineered systems, where low but sustained fluxes maintain biological activity in the absence of bulk supply. This reframing has direct implications for material selection. Membranes that balance ionic conductivity with moderate permeability—rather than maximizing selectivity—may enable operation in regimes where metabolic activity is sustained without excessive substrate accumulation. Consequently, separator design becomes a problem of transport tuning, in which permeability is adjusted to match biological demand rather than minimized unconditionally.

### 5.3. Current-coupled transport and intrinsic stabilization

The inclusion of current-dependent transport introduces an additional layer of coupling between electrochemical and biological processes. Because solvent drag scales with current density, variations in metabolic activity can influence the net substrate flux through the separator. This coupling generates an intrinsic feedback mechanism in which increased current output reduces net substrate delivery, thereby moderating further increases in metabolic activity. Conversely, reduced activity weakens this opposing transport contribution, allowing diffusive flux to dominate. Although secondary to diffusion under most conditions, this feedback may contribute to the stability of the balanced regime by damping fluctuations in substrate availability. Such self-regulating behavior is characteristic of coupled transport systems and suggests that electrochemical operation can indirectly influence mass transfer in a manner that promotes dynamic stability.

### 5.4. Sensitivity, robustness, and parameter uncertainty

A key feature of the flux-based formulation is its relative robustness to parameter uncertainty. Because regime classification depends on the ratio between transport capacity and metabolic demand, moderate variations in individual parameters—such as separator permeability, catholyte concentration, or maintenance flux—do not necessarily alter the qualitative system behavior. This robustness is particularly important given the variability inherent to bioelectrochemical systems. Effective diffusivity depends on membrane microstructure and hydration, while maintenance demand varies with microbial community composition and physiological state. Despite these uncertainties, systems designed near the balanced regime are expected to retain stable operation over a broad parameter space. This property enhances the practical relevance of the framework, as it reduces the sensitivity of design criteria to poorly constrained variables.

### 5.5. Practical implications for reactor and material design

The theoretical framework leads to several concrete implications for the design of autonomous bioelectrochemical systems: (i) Permeability as a design variable. Separator permeability should be tuned to achieve a balance between transport supply and metabolic demand, rather than minimized to suppress crossover. This shifts the design objective from selectivity to controlled transport. (ii) Catholyte as an engineered reservoir. The catholyte compartment can be deliberately configured to store metabolically accessible substrates, enabling sustained operation through passive delivery. (iii) Coupled parameter optimization. Separator thickness, effective diffusivity, substrate concentration, and operating current density must be co-optimized, as they jointly determine the transport–reaction balance. (iv) Autonomy- oriented optimization. For applications prioritizing long-term stability, design strategies should favor operation in the balanced regime, even at the expense of peak coulombic efficiency. These considerations define a design space distinct from that of conventional high-performance MFCs, emphasizing resilience and longevity over maximum instantaneous output.

### 5.6. Extension to microbial electrolysis cells

The transport–reaction framework is directly extensible to microbial electrolysis cells, in which externally applied voltage drives higher current densities and alters transport conditions [102, 103]. In these systems, the absence of cathodic oxygen and the presence of stronger ionic fluxes modify the relative contributions of diffusive and current-coupled transport [104]. Despite these differences, the same fundamental balance between substrate supply and maintenance demand governs anodic biofilm stability. Order-of-magnitude analysis indicates that, due to higher effective permeabilities and current densities, transport fluxes in such systems can exceed maintenance requirements by several orders of magnitude [102, 105]. As a result, operation may naturally shift toward supply-dominated regimes unless permeability is carefully controlled. This extension highlights the generality of the framework and suggests that the same dimensionless criterion can be used to guide the design of both electricity-generating and hydrogen-producing bioelectrochemical systems.

### 5.7. Limitations of the present framework

The analysis is based on a simplified representation of transport and biological processes and is therefore subject to several limitations. First, the model assumes constant transport coefficients and neglects spatial heterogeneity within both the membrane and the biofilm. In practice, variations in hydration, fouling, or biofilm structure may alter effective transport properties [106]. Second, microbial metabolism is represented through a constant maintenance flux, whereas real biofilms exhibit dynamic behavior, including adaptation, decay, and shifts in metabolic pathways during prolonged starvation [107]. Third, the framework does not explicitly account for competing processes such as oxygen crossover, alternative electron acceptors, or microbial interactions (e.g., methanogenesis), which may influence electron recovery and substrate utilization [53, 108]. Finally, pH gradients and ionic transport are not explicitly coupled to substrate transport, although they can significantly affect both membrane properties and microbial kinetics [109]. While these simplifications limit quantitative prediction under complex conditions, they do not invalidate the central insight that system behavior is governed by a balance between transport capacity and maintenance demand.

### 5.8. Broader theoretical implications

The framework developed in this work extends classical reaction–transport theory to a regime in which maintenance metabolism rather than growth dominates system dynamics. This shift introduces a new class of transport-limited systems characterized by low reaction rates and long characteristic timescales. Within this context, membrane transport emerges not only as a constraint but as a control mechanism capable of stabilizing biological activity. The resulting conceptual model bridges electrochemical engineering, microbial kinetics, and membrane science, providing a unified basis for analyzing bioelectrochemical systems under resource-limited conditions. More broadly, the results suggest that similar transport–reaction balances may govern other systems in which biological activity is sustained by passive fluxes, including membrane bioreactors, natural sediments, and low-energy ecological niches.

### 5.9. Testable predictions and validation pathways

The theory yields several experimentally testable predictions:

- Systems designed near the balanced regime should sustain measurable current during extended periods without external substrate input.
- Increasing catholyte substrate concentration or reservoir volume should extend operational lifetime in proportion to available substrate.
- Variations in current density should produce measurable changes in net substrate flux due to current-coupled transport effects.

Validation of these predictions through controlled experiments would provide strong support for the proposed framework and enable refinement of model parameters for specific materials and reactor configurations.

### 5.10. Synthesis

Taken together, the results establish a coherent interpretation of substrate crossover as a controllable transport process capable of sustaining microbial metabolism under substrate-limited conditions. By linking membrane properties, operating conditions, and biological demand within a unified framework, the analysis defines a new regime of operation in which passive transport enables long-term stability. This perspective expands the design space of bioelectrochemical systems and provides a theoretical foundation for the development of autonomous reactors optimized for resilience, low maintenance, and sustained functionality.

## 6. Conclusion

This work establishes a rigorous transport–reaction framework that re- defines substrate crossover in dual-chamber microbial fuel cells (MFCs) as a controllable mechanism for sustaining microbial activity under substrate-limited conditions. By explicitly coupling membrane-mediated transport with biofilm maintenance metabolism, the analysis demonstrates that long-term system viability is governed by the balance between passive substrate flux and minimal metabolic demand. The introduction of the maintenance crossover Damköhler number provides a dimensionless and hydrodynamics-independent criterion for regime classification. This formulation enables clear identification of transport-limited, balanced, and crossover-dominated regimes, and shows that autonomous operation without continuous feeding is theoretically attainable when transport capacity is tuned to maintenance requirements. Beyond its theoretical contribution, the framework implies a fundamental shift in MFC design philosophy. Rather than minimizing membrane permeability to suppress crossover, optimal performance in autonomy-oriented systems requires controlled permeability that enables sustained, low-level substrate delivery. In this context, the separator becomes a tunable transport interface, while the catholyte acts as an engineered substrate reservoir supporting long-term biofilm viability. Although based on idealized assumptions, the dimensionless structure of the model ensures generality and provides a scalable basis for experimental validation and multiphysics extension. Future work should focus on quantifying transport parameters in real materials and verifying regime behavior under controlled conditions. Overall, this study expands the design space of bioelectrochemical systems by demonstrating that passive transport can be harnessed to achieve resilience and long-term autonomous operation.

